# Distinct cis-acting elements combinatorically mediate co-localization of mRNAs encoding for co-translational interactors in cytoplasmic clusters in *S. cerevisiae*

**DOI:** 10.1101/2025.02.19.639054

**Authors:** Qamar Shhade, Marianna Estrada, Rawad Hanna, Muhammad Makhzumy, Hagit Bar-Yosef, Guenter Kramer, Bernd Bukau, Ayala Shiber

## Abstract

Many newly synthesized proteins assemble co-translationally, providing a vital mechanism to prevent subunit misfolding in the crowded cytoplasm. Initial evidence indicates that the spatial organization of mRNAs aids this assembly, but it is unclear how these mRNAs are organized and how common this mechanism is. We used single-molecule Fluorescence *in situ* Hybridization in *Saccharomyces cerevisiae* to examine the spatial organization of mRNAs encoding subunits of various cytosolic complexes involved in critical cellular functions, such as fatty acid synthesis, glycolysis, translation and various amino acid biosynthesis. We found that mRNAs of the same protein complex often co-localize in specific cytoplasmic clusters. Additionally, we observed that the mRNAs encoding enzymes of biosynthetic pathways are organized in cytosolic clusters. Focusing on mRNAs encoding fatty acid synthase complex subunits, we discovered that non-coding cis elements significantly influence mRNA localization in an additive manner. Specifically, 5’ and 3’ UTRs, together with further upstream and downstream regions, facilitate co-localization. Inhibiting mRNA co-localization impaired growth when complex activity was essential, highlighting the importance of mRNA spatial organization for cellular survival. Transiently disrupting mRNA translation also affected clustering, indicating that both the nascent chains and mRNA sequence targeting cues are combinatorically contributing to spatial organization. Proteomics analysis demonstrates the impact of cis-elements on the abundance of the encoded subunits, as well as the entire pathway. In summary, we provide evidence that mRNA co-localization in cytoplasmic foci is coordinated by complementary mechanisms crucial for co-translational assembly, allowing efficient regulation of protein complex formation and entire pathways.

## Introduction

The majority of cellular proteins can only function as parts of multi-subunit complexes, working together to achieve coordinated functions. This organization is highly conserved across species (Krogan et al., 2006; Reid et al., 2010). The assembly of protein complexes is a highly intricate process that ensures proper folding and functional integrity. A crucial element is co-translational interactions, where subunits start assembling in coordination with the translation process. Co-translational assembly is a well-established phenomenon in eukaryotes, conserved from yeast to humans (Bertolini et al., 2021; Duncan & Mata, 2011; Halbach et al., 2009; Hampoelz et al., 2019; Kamenova et al., 2019; Khan et al., 2022; Lautier et al., 2021; Mallik et al., 2025; Seidel et al., 2022, 2023; Shiber et al., 2018; Venezian et al., 2024). This process is initiated as the polypeptide chain emerges from the ribosome exit tunnel. Specific interface residues, termed “hotspots” can initiate assembly interactions through high-energy binding, anchoring interface formation early in the translation process (Venezian et al., 2024). This mechanism ensures timely formation of functional complexes and protects nascent polypeptide chains from misfolding and aggregation (Boguta, 2022; Khan & Fox, 2023). Assembly interactions are coordinated with ribosome-associated molecular chaperones, such as the Ribosome-Associated Complex (RAC), which safeguard nascent chains from premature or off-pathway interactions. This coordination ensures proper folding and assembly, preserving proteome organization and cellular homeostasis (Shiber et al., 2018).

This raises the question: Are mRNAs spatially organized within the cytosol to facilitate co-translational assembly interactions? The spatial organization of mRNAs has been demonstrated to play a crucial role in the regulation of gene expression in eukaryotes (Cassella & Ephrussi, 2022; Das et al., 2021; Horste et al., 2023; Mikl et al., 2022; Weatheritt et al., 2014). While prokaryotes utilize operons for co-localized synthesis of complex subunits and functionally related genes (Nair et al., 2021; Shieh et al., 2015), eukaryotes employ different strategies for protein synthesis coordination.

Localization of specific mRNAs at subcellular sites is common in eukaryotes. For example, mRNAs encoding for endoplasmic reticulum (ER) and mitochondrial proteins are targeted to the respective organellar membranes (Cohen et al., 2022; Eliyahu et al., 2012; Gadir et al., 2011; Garcia Arceo, 2023; Gerst, 2008; Zabezhinsky et al., 2017). Membrane targeting is facilitated by mechanisms like the Signal Recognition Particle (SRP) pathway (Hermesh & Jansen, 2013; Pyhtila et al., 2008) and RNA-binding proteins that regulate mRNA localization (Lesnik et al., 2015).

In various tissues, subcellular mRNA localization involves active transport by molecular motor proteins and adaptors along cytoskeletal tracks, and localized mRNA degradation (Cassella & Ephrussi, 2022). In neurons, cytosolic mRNAs encoding the Cox7c (Cytochrome C Oxidase Subunit 7C) mitochondrial protein are co-transported with mitochondria along axons, enabling localized translation (Cohen et al., 2022; Eliyahu et al., 2012). During early Drosophila embryonic development, specific mRNAs localize to the oocytés poles, establishing morphogen gradients (Gagnon & Mowry, 2011; Johnstone & Lasko, 2003; Lehmann’ & Nüsslein-Volhard’, 1986). Even single cell organisms like yeast exhibit active mRNA transport, such as the transport of *ASH1* (*Asymmetric Synthesis of HO*) mRNA to the bud tip of a dividing cell for mating-type switching (Paquin & Chartrand, 2008). Furthermore, mRNAs encoding several mating pathway factors, including *ASH1*, have been found to co-localize at the ER, in both growing and pheromone-stimulated yeast cells, thereby modulating pheromone sensitivity (Nair et al., 2021). Together, these examples highlight the diverse roles of mRNA co-localization in cellular functions and spatially regulated gene expression.

Another type of co-localization of mRNAs is induced by stress conditions, such as starvation and heat shock, leading to co-localization of mRNAs and RNA-binding proteins in stress granules (SGs). SGs provide a protective environment for transcripts and ensure coordinated translation upon stress recovery. Under glucose deprivation, for example, mRNAs encoding human regulatory subunits Rpt1 and Rpt2 of the proteasome co-localize at distinct subcellular sites. This process depends on the CCR4-NOT1 complex, which regulates multiple aspects of the mRNA lifecycle, including its synthesis, translation, and degradation (Panasenko et al., 2019). In growing yeast cells, specific glycolytic mRNAs localize to granules, which recruit mRNA decay factors to become P-bodies during glucose starvation, suggesting an interplay between mRNA translation and degradation (Balagopal & Parker, 2009; Buchan & Parker, 2009; Haimovich et al., 2016; Hoyle & Ashe, 2008; Lui et al., 2014). Notably, these granules were recently found to represent a unique type of glycolytic mRNA-protein granules, termed Core Fermentation (CoFe) granules, in which coordinated localization and translation regulate glucose fermentation under various conditions (Morales-Polanco et al., 2021). Moreover, granules containing mRNAs encoding for translation factors are contributing to budding and polarized growth, proposing a model where specific mRNAs are directed to granules for localized protein synthesis (Pizzinga et al., 2019).

Several studies have examined the significance of mRNA co-localization in protein complex assembly. Early analysis of sarcomere pattern establishment in muscle cells, identified coordinated expression and assembly of myosin, titin, and other proteins (Fulton & L’ecuyer, 1993). In migrating fibroblasts, mRNA localization to the leading protrusions is crucial for the Arp2/3 (Actin Related Protein 2/3) complex assembly, influenced by extracellular stimuli and cytoskeletal components (Mingle et al., 2005). mRNA localization together with nucleoporins condensation is also critical for nuclear pore complex (NPC) assembly, with co-translational assembly of nuclear pore subcomplexes ensuring ordered assembly of the highly intricate nuclear pore complex (Hampoelz et al., 2019; Lautier et al., 2021; Penzo & Palancade, 2023). The biogenesis of the mammalian TFIID multi-subunit complex follows a hierarchical assembly, driven by co-translational dimerization of several subunits. Endogenous TFIID subunits have been observed in close proximity to *TAF1* mRNA foci within the cytoplasm of human cells (Bernardini et al., 2023; Bernardini & Tora, 2024; Kamenova et al., 2019). Mallik et al. recently performed a broad investigation into the structural determinants of co-translational protein complex assembly across bacteria, yeast, and humans. By utilizing AlphaFold-predicted interface structures, they found that subunits with highly intertwined interfaces are more likely to assemble co-translationally. Their analysis of several yeast complexes revealed active co-translational assembly in multiple cases, with two of these highly large multi-subunit complexes exhibiting pronounced mRNA co-localization. However, the study also finds that other co-translationally assembling complexes displayed only minimal mRNA co-localization, and the reasons for this variability remain unclear (Mallik et al., 2025). Similarly, the ATAC (ADA-two-A-containing) and SAGA (Spt-Ada-Gcn5-acetyltransferase) transcriptional co-activator mega complexes both employ co-translational assembly mechanisms; however, they differ in their cellular distribution of protein subunits as well as encoding transcripts (Yayli et al., 2023).

Taken together, these sporadic and conflicting examples bring forward the key questions: How common is the phenomenon where mRNAs are spatially organized within the cytosol to facilitate co-translational assembly interactions? What features regulate this spatial organization? How does this arrangement contribute to the efficiency of protein complex assembly, functionality and cellular viability?. This study investigates the spatial distribution of mRNAs coding for protein complex subunits and metabolic pathways, focusing on the role of localized translation in *S.* cerevisiae, across various cytoplasmic processes. We identify mechanisms supporting mRNA co-localization and examine the impacts of failed mRNA localization on protein assembly and functionality. Our research shows that mRNAs encoding for different subunits of the same protein complex often cluster in cytoplasmic foci. By swapping sequence elements, we identify specific upstream and downstream elements affecting mRNA co-localization and complex formation, suggesting spatial mRNA organization is a key regulatory mechanism in protein complex assembly. The co-localized synthesis of protein subunits may be essential for maintaining a functional proteome, especially given the instability of orphan subunits.

## Results

### Co-localization of mRNAs encoding protein complex subunits in diverse pathways

To investigate whether co-translational assembly interactions are facilitated by localized translation across various cytoplasmic pathways, we analyzed the *in vivo* localization of mRNAs encoding complex’ subunits using single-molecule fluorescence in situ hybridization (smFISH) in yeast (Zenklusen et al., 2008). They were selected to encompass a range of cellular functions, structural types and regulatory mechanisms, including fatty acid synthesis (fatty acid synthase, FAS), glycolysis (Phosphofructokinase, PFK), translation (Translation Initiation Factor eIF2 and the <u>a</u>mino<u>a</u>cyl-tRNA synthetases complex), and amino acids biosynthesis - tryptophan and Arginine pathways (Supplementary Table 1). We selected complexes where we previously identified co-translational assembly mode (Shiber et al., 2018). As negative controls, we assessed co-localization of *ASH1* mRNA at the bud neck and cytoplasmic *FAS1* mRNA, as well as co-localization of cytoplasmic mRNAs encoding subunits of different complexes, such as *TRP2* and *MES1 (TRyPtophan2 & MEthionyl-tRNA Synthetase)*. For a positive control for maximum co-localization detectable under our experimental conditions, we labeled the same transcript, *FAS1,* with two different fluorescent probes.

The smFISH analysis of endogenous mRNA distribution (Figure 1) revealed consistent mRNA co-localization patterns for transcripts encoding different subunits of the same protein complexes. In contrast, negative control pairs *ASH1* and *FAS1*, as well as *TRP2* and *MES1*, showed very low levels of co-localization. The complexes analyzed showed a range of co-localization levels, spanning from 11% to 54%, yet all showed significance co-localization levels in compared to the two negative controls. *FAS1* and *FAS2* mRNAs, encoding the α and β subunits of fatty acid synthase, exhibited the highest level of co-localization among the studied complexes: a median of 54% mRNA co-localization per cell. Similarly, *PFK1* and *PFK2*, encoding subunits of phosphofructokinase, exhibited 50% co-localization. *GUS1* and *ARC1*, encoding components of the aminoacyl-tRNA synthetases complex, also displayed high levels of co-localization of ∼44%. *GUS1* and *MES1* of the same complex displayed ∼27%, potentially indicating an underlying assembly hierarchy. Interestingly, selective ribosome profiling analysis of this complex co-translational assembly mechanism (Shiber et al., 2018) also suggested a similar hierarchy. *TRP2* and *TRP3* transcripts, encoding components of the Anthranilate synthase in the Tryptophan biosynthetic pathway displayed 29% co-localization levels. *SUI2* and *GCD11* transcripts (*SUppressor of Initiator codon 2 & General Control Derepressed 11*) encoding subunits of the eIF2 complex (eukaryotic translation initiation factor 2) showed 27% while *SUI2* and *SUI3* mRNA encoding subunits of the same complex showed 11%, potentially indicating an underlying assembly hierarchy, as suggested for the aminoacyl-tRNA synthetases complex.

**Figure 1.**
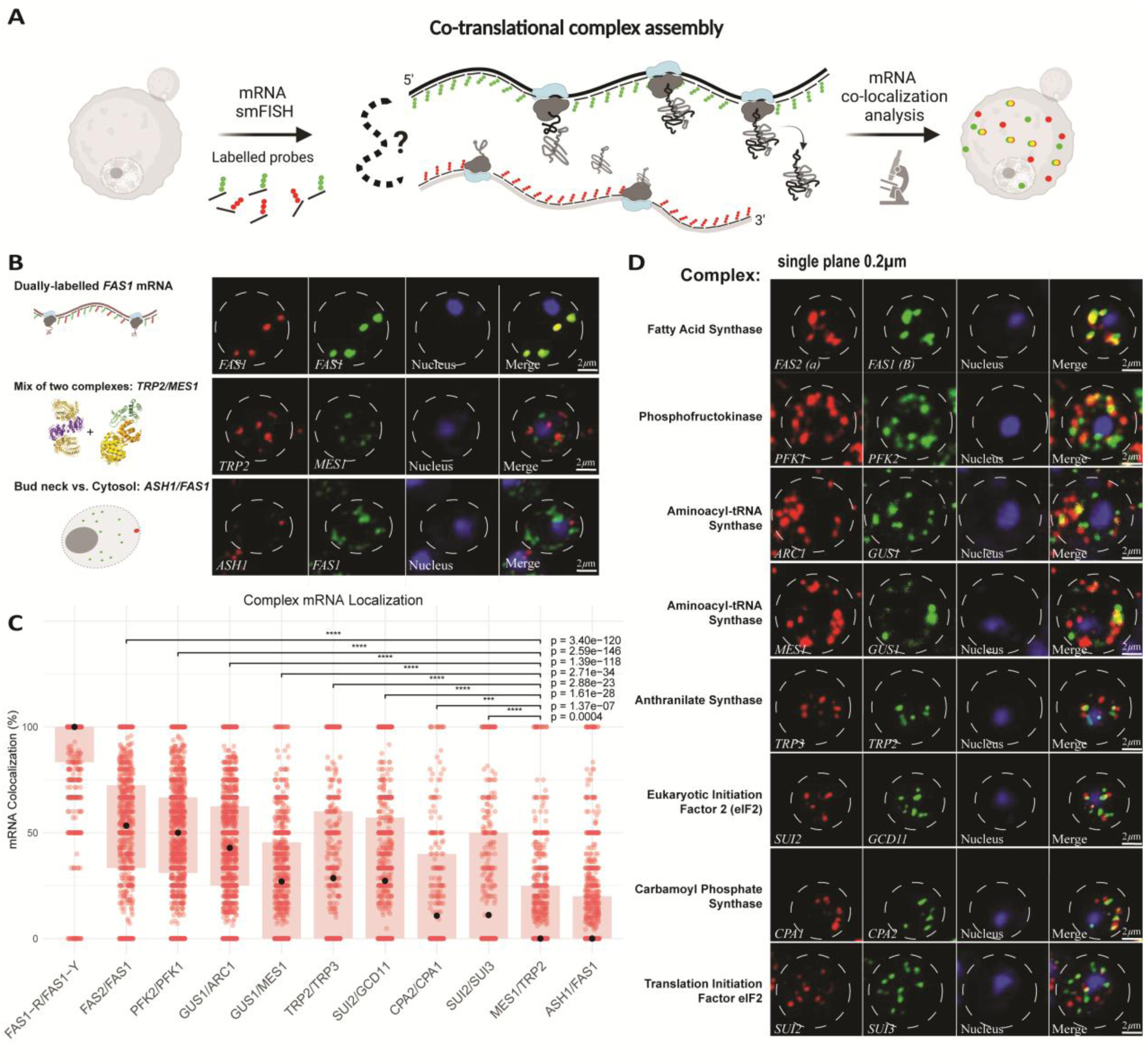
Colocalization of mRNAs encoding complex subunits in diverse pathways. **A.** Schematic workflow illustrating the smFISH methodology used to analyze the co-localization of mRNAs encoding subunits of a single protein complex. **B.** Control Experiments for smFISH Detection. Single z-plane images showing smFISH detection of control conditions in wild-type yeast strains. Positive control: probes of different colors binding the same mRNA. Specificity controls: a mix of unrelated mRNAs (*TRP2*, *MES1*) and (*ASH1*, *FAS1)*. Yeast cells were grown to log phase, fixed with formaldehyde, and hybridized with smFISH probes in two channels: channel one (red) and channel two (green). Nuclei were stained with Hoechst (blue) for visualization. Scale bar = 2 µm. **C.** Quantification of mRNA co-localization: Boxplot showing the proportion of co-localized smFISH-detected mRNAs per cell was analyzed. Co-localization was assessed pairwise for dually labeled mRNAs encoding subunits of a single protein complex, functionally unrelated mRNAs, and non-co-localized mRNAs. The box represents the interquartile range, with the median indicated by a point. Each data points represents an individual cell (n > 1036 cells per detection). At least two independent experiments were conducted. Statistical analysis revealed highly significant p-values compared to *TRP2*/*MES1* negative controls. **D.** Single z-plane images showing smFISH detection of the indicated mRNAs in wild-type yeast strains. Yeast cells were grown to log phase, fixed with formaldehyde, and hybridized with smFISH probes for mRNA detection in two channels: channel one (red) and channel two (green). Nuclei were stained with Hoechst (blue) for visualization. Scale bar = 2 µm.

*CPA1* and *CPA2* (*Carbamyl Phosphate synthetase A* 1,2) of the Arginine pathway also displayed 11% co-localization levels. These results indicate that co-localization of mRNAs encoding different subunits of the same protein complex is widespread. This organization may facilitate co-translational protein-protein interactions to promote proper folding in the crowded cytoplasm and protecting nascent chains from misfolding. Furthermore, the observed variation in co-localization levels between complexes suggests the existence of assembly hierarchies and / or tight regulatory mechanisms, which may selectively permit complex formation only under certain conditions.

### Co-localization of mRNAs encoding interacting proteins within biosynthetic pathways

We expanded our analysis to mRNAs encoding enzymes of three major metabolic pathways: fatty acid synthesis, arginine biosynthesis, and glycolysis (Supplementary Table 2). We previously observed that components of these pathways exhibited transient co-translational interactions of their nascent polypeptides (Shiber et al., 2018), including components of distinct enzymes within a pathway, indicating that co-localized translation may contribute to pathway regulation.

Supporting our model, mRNAs encoding co-translational protein interactors within the same pathway often show co-localization in cytosolic foci (Figure 2). For example, *FAS1* and *ACC1* mRNAs, which encode Acetyl-CoA Carboxylase (Acc1p) and the β subunit of the FAS complex, catalyze consecutive reactions in lipid synthesis and have been found to transiently interact co-translationally. Our smFISH analysis revealed that *ACC1* and *FAS1* mRNAs are significantly close to each other (Figure 2. C), with a median of 58% co-localized mRNAs per cell. Similarly, *PFK2* and *PYC2* mRNAs exhibit notable co-localization (Figure 2). *PFK2* encodes phosphofructokinase 2, a key glycolytic enzyme, while *PYC2* encodes pyruvate carboxylase, which converts pyruvate to oxaloacetate in gluconeogenesis. Despite their involvement in distinct, yet interconnected metabolic pathways, these proteins are functionally coordinated in cellular energy metabolism. smFISH analysis showed that *PFK2* and *PYC2* mRNAs are in close proximity, with a median of 50% co-localized mRNAs per cell. In contrast, the control pairs *ASH1* and *FAS1,* as well as *TRP2* and *MES1*, showed no significant co-localization.

**Figure 2.**
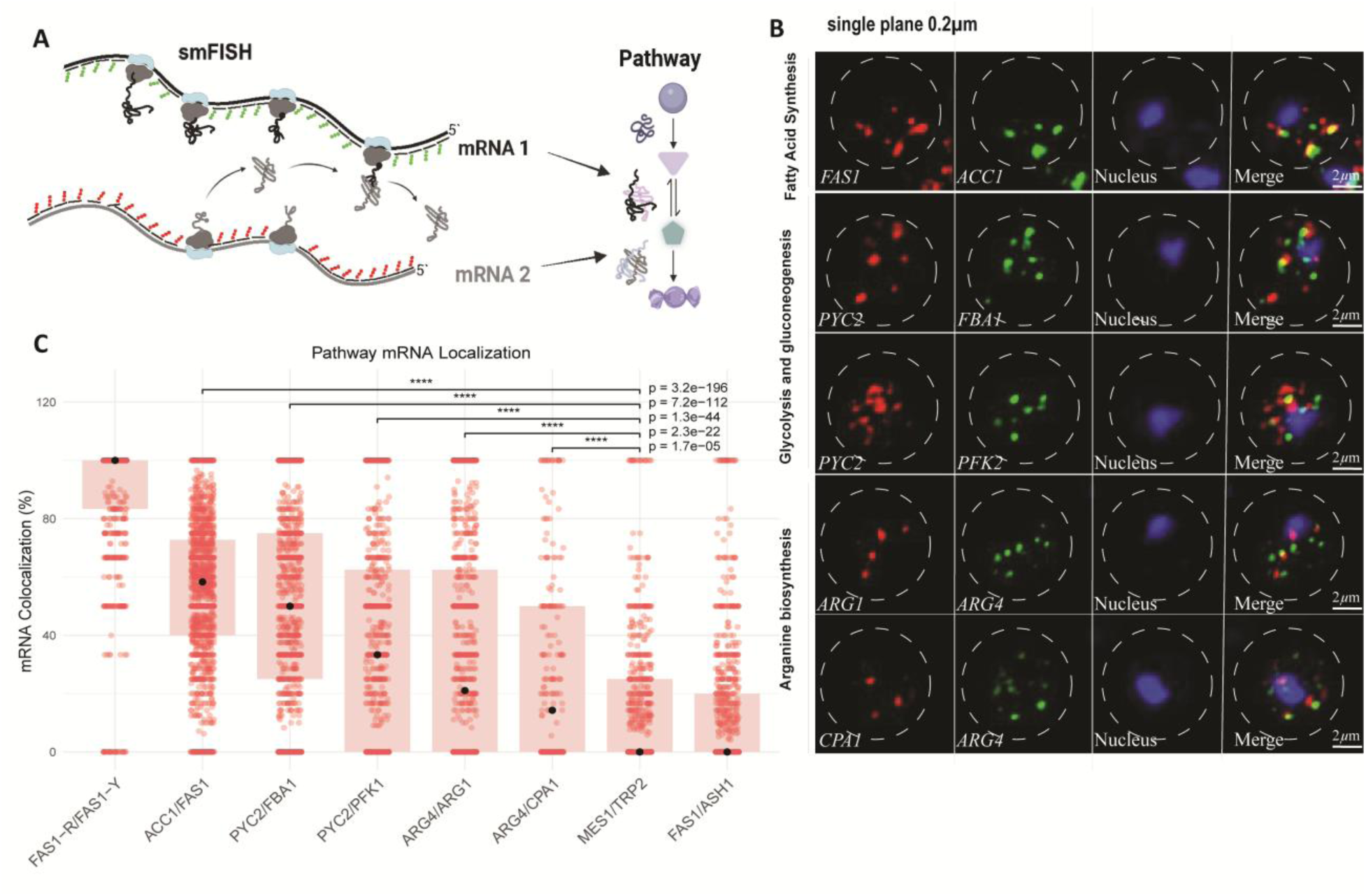
Colocalization of mRNAs encoding interacting proteins of biosynthetic pathways. **A**. Schematic workflow illustrating the smFISH methodology used to analyze the co-localization of mRNAs encoding proteins within a single biosynthetic pathway. **B.** Single z-plane images showing smFISH detection of the indicated mRNAs in wild-type yeast strains. Yeast cells were grown to log phase, fixed with formaldehyde, and hybridized with smFISH probes for mRNA detection in two channels: channel one (red) and channel two (green). Nuclei were stained with Hoechst (blue) for visualization. Scale bar = 2 µm. **C.** Quantification of mRNA co-localization: Boxplot showing the proportion of co-localized smFISH-detected mRNAs per cell. Co-localization was assessed pairwise for dually labeled mRNAs encoding interactors within a single biosynthetic pathway, functionally unrelated mRNAs, and non-co-localized mRNAs. The box represents the interquartile range, the median indicated by a black dot. Each data point represents an individual cell (n > 778 cells per detection). At least two independent experiments were conducted. Statistical analysis revealed highly significant p-values compared to *TRP2*/*MES1* negative control.

Our findings connect mRNA co-localization with co-translational interactions of proteins in functional protein networks, potentially coordinating gene expression at the translational level. The regulation of translation, including the formation of mRNA-containing SG condensates, is as a critical mechanism responsive to nutrient availability, pheromones, stress, and pathogen infection (Leprivier et al., 2013; Levy et al., 2024; Morales-Polanco et al., 2021; Musa et al., 2016; Nair et al., 2021). Our results suggest that even under optimal growth conditions with abundant nutrients, cells fine-tune various pathways’ functions by regulating the spatial organization of mRNAs encoding key metabolic enzymes.

### Co-localization of mRNAs occurs in the cytosol

We examined whether co-localization of functionally related mRNAs begins in the nucleus, close to their transcription sites (Supplementary Table 3), or later in the cytoplasm where translation occurs. Analysis of localization patterns in different compartments revealed that mRNAs encoding co-translational interactors primarily co-localize in the cytosol (Figure 3), with co-localization levels ranging from 11% to 55%, while nuclear mRNA showed a median of 0% co-localization. Notably, we detected high nuclear co-localization for dually labeled *FAS1* mRNAs, indicating that all probes (*FAS1*-Red and *FAS1-*Yellow) target the same transcription site. These results suggest that mRNAs come together in the cytosol rather than in the nucleus, indicating cytoplasmic mechanisms driving co-localization.

**Figure 3:**
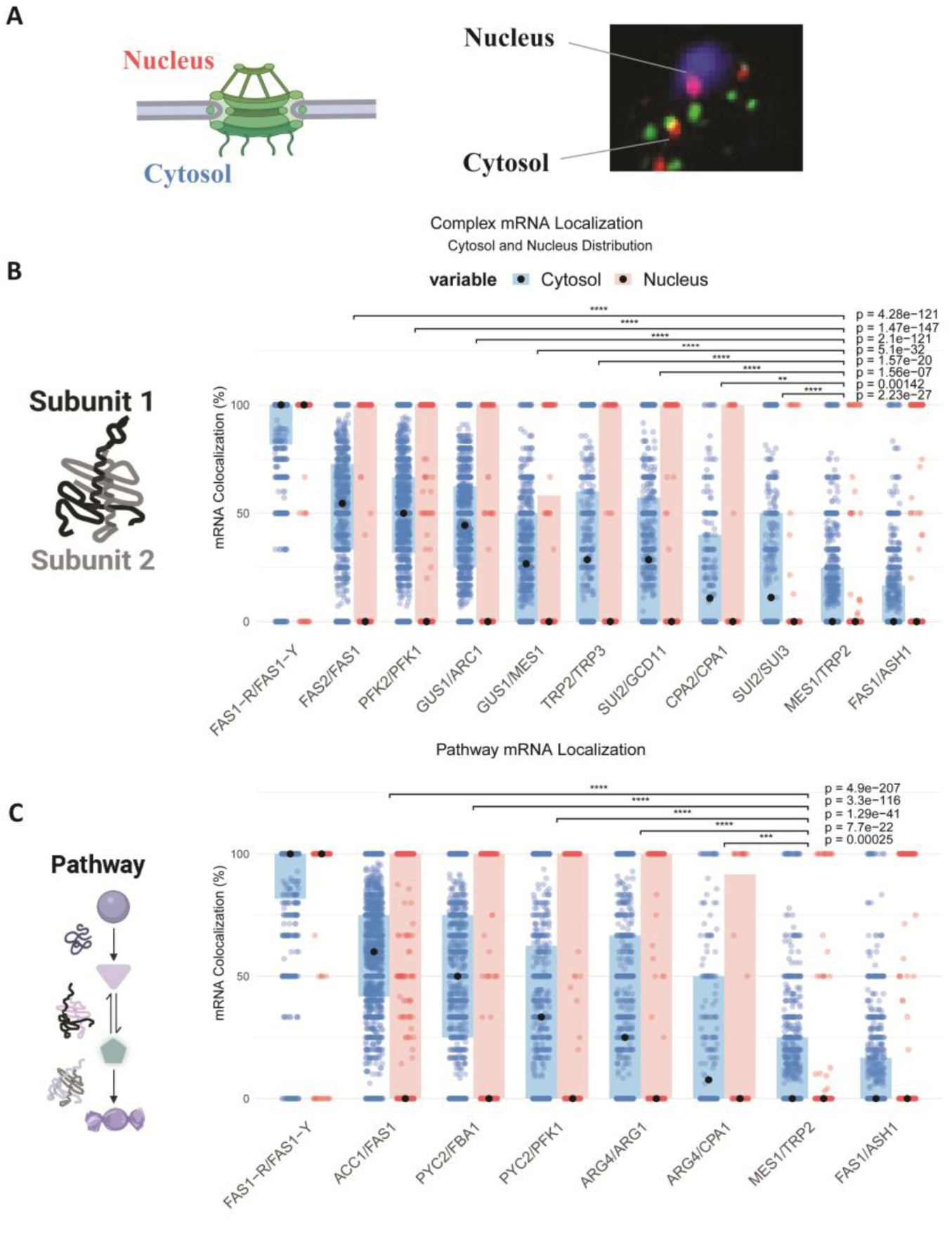
Colocalization of mRNAs encoding for co-translational interactors occurs in the cytoplasm. **A.** Schematic representation of mRNA localization in the cytoplasm (blue) versus nuclei (light pink): A diagram illustrating the spatial distribution of mRNAs within yeast cells, highlighting mRNA localization in the cytoplasm versus nuclei. **B.** Quantification of mRNA co-localization in the cytoplasm and nuclei for subunits of a single protein complex: Boxplot showing the proportion of co-localized mRNAs per cell detected using smFISH probes. Co-localization was analyzed for mRNAs encoding subunits of a single protein complex in the cytoplasm and nuclei. **C.** Quantification of mRNA co-localization in the cytoplasm and nuclei for biosynthetic pathway proteins: Boxplot depicting the proportion of co-localized mRNAs per cell for mRNAs encoding proteins within a single biosynthetic pathway. Detection was carried out separately in the cytoplasm and nuclei. Data are presented as boxplots, with the box representing the interquartile range and the black dot indicating the median. Each data point corresponds to an individual cell (n > 778 cells per detection). The results are based on at least two independent experiments, showing highly significant p-values compared to negative control (*TRP2*/*MES1*).

### mRNA clusters in cytoplasmic foci dedicated to complex assembly

We investigated how many mRNA molecules are present in each cytosolic assembly focus. smFISH’s remarkable sensitivity allows detection of single mRNA molecules and estimation of mRNA levels based on foci fluorescence intensity (Zenklusen et al., 2008). To assess mRNA co-localization within distinct foci, we classified foci into two groups: those with less than 2 mRNAs (<2) and those with 2 or more mRNAs (≥2). Analysis showed that multi-mRNA foci had significantly higher co-localization rates across most mRNA pairs (Figure 4). For example, *FAS1* and *FAS2* co-localized in 75% of multi-mRNA foci, *PFK*1-*PFK2* in 60%, and *GUS1*-*ARC1* in 53.7%. Similarly, analysis of mRNAs encoding different subunits of the same biosynthetic pathways revealed trends such as 66.6% co-localization for *ACC1/FAS1*. These findings suggest that mRNA spatial organization in the cytoplasm is coordinated, with multiple mRNAs brought to specific subcellular foci to facilitate assembly and biosynthetic pathways formation.

**Figure 4.**
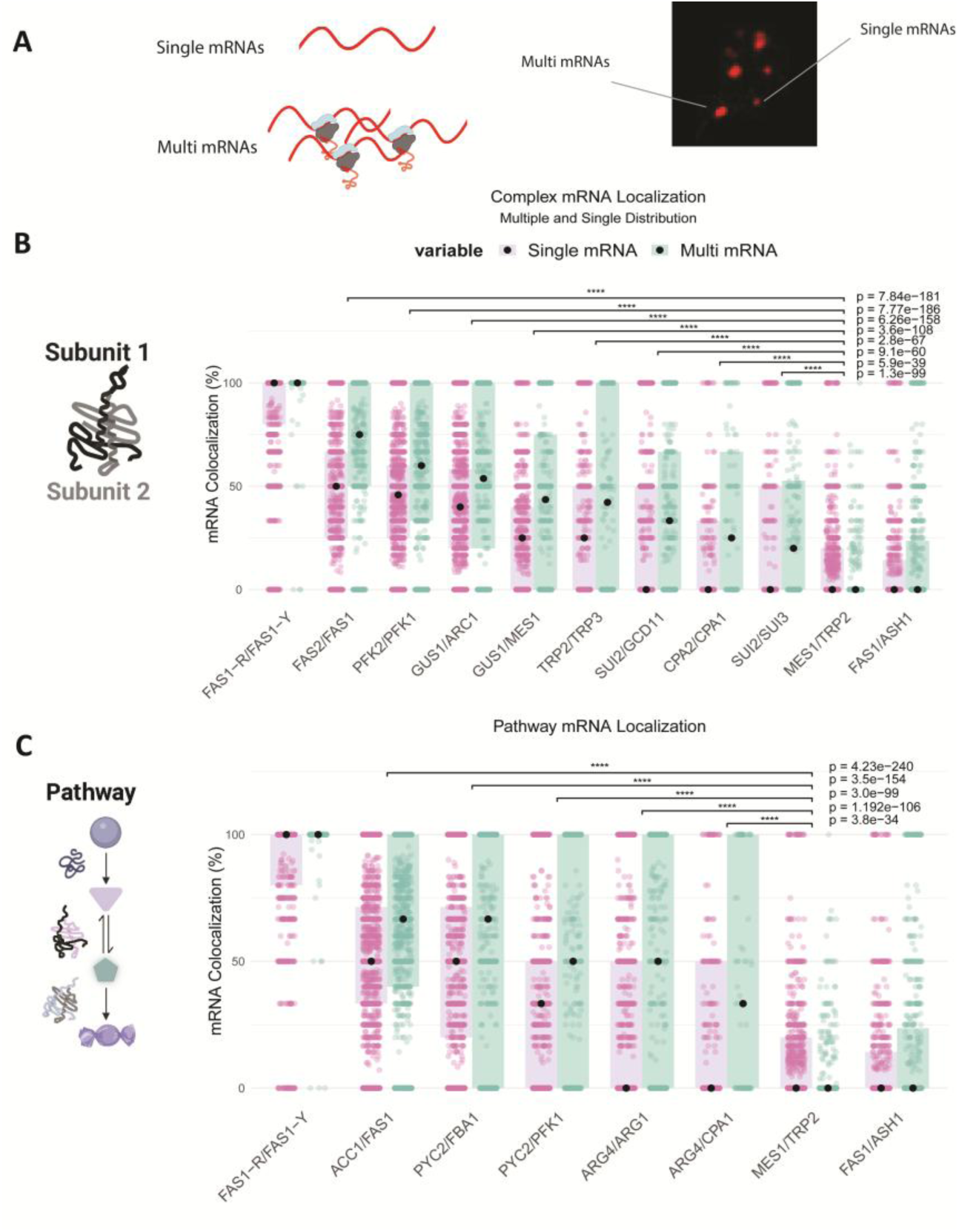
Co-translational interactors’ mRNA clusters in cytoplasmic foci: **A.** Schematic representation of single (purple) and multi-mRNA (green) foci: Diagram illustrating the spatial organization of mRNAs within cytoplasmic foci. Single mRNA foci represent individual mRNAs localized as discrete spots, while multi-mRNA clusters consist of two or more mRNAs grouped together in a single focus. smFISH was performed on wild-type yeast cells, targeting the coding regions of specific mRNAs of interest. **B.** Quantification of mRNA clustering for subunits of a protein complex: Boxplot showing the proportion of co-localized mRNAs per cell, categorized by the number of mRNAs per focus. Foci with fewer than two mRNAs (<2) were compared to multi-mRNA clusters containing two or more mRNAs (≥2). Data are presented as boxplots, with the box representing the interquartile range and the black dot indicating the median. Each data point represents an individual cell. Results from at least two independent experiments indicate highly significant p-values compared to negative control (*TRP2*/*MES1)*. **C**. Quantification of mRNA clustering for proteins of a biosynthetic pathway: Analyzed similarly to panel B, showing the proportion of co-localized mRNAs per cell for mRNAs encoding proteins within a single biosynthetic pathway. same boxplot format as in panel B.

### Translation-dependent co-localization of mRNAs encoding various complexes’ subunits

We investigated how inhibition of translation affects co-localization of mRNAs encoding for FAS, PFK, TRP, CPA, eIF2, and aminoacyl-tRNA synthetase complexes’ subunits. The cells were treated with Puromycin, as it incorporates into the C-termini of elongating nascent chains, blocking further extension, resulting in premature termination and nascent chains release (Baliga et al., 1970; Bresson et al., 2020; Semenkov et al., 1992, 1992; Wu, 2003). To ensure Puromycin effectiveness, cells were grown in the presence of SDS and proline, sensitizing the cells. Polysome analysis demonstrates Puromycin treatment significantly reduced translation levels resulting in decreased polysomes levels (Figure 5B).

**Figure 5.**
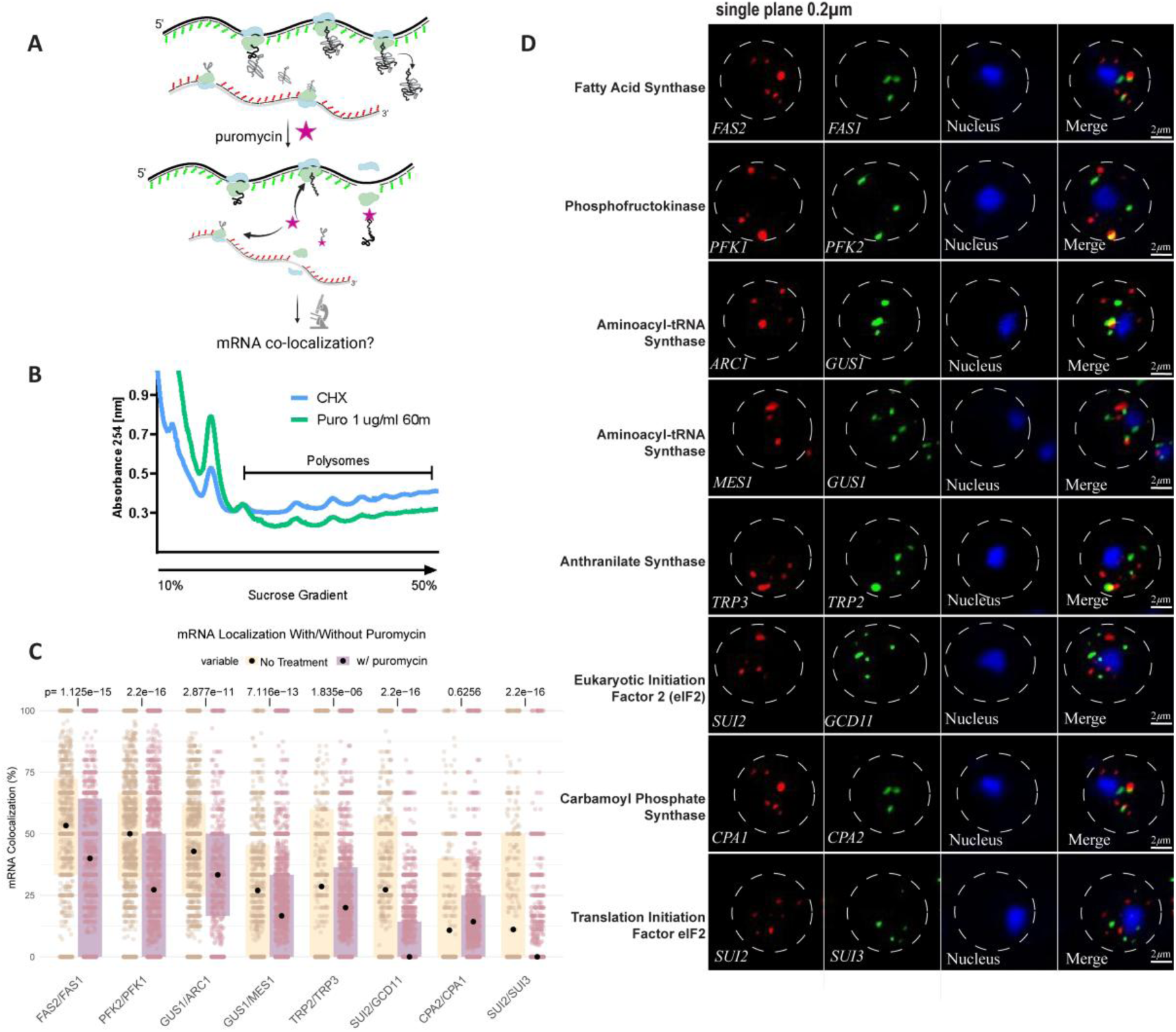
Translation-dependent and independent co-localization of mRNAs encoding various complexes’ subunits. **A.** Schematic illustration of puromycin action on ribosome–nascent chain complexes. Puromycin incorporation causes premature termination and release of nascent chains, allowing analysis of translation-dependent contributions to mRNA co-localization. **B.** Polysome profiling of WT cells treated with puromycin (1 µg/ml, 60 min) compared to cycloheximide (CHX) control. Ribosome run-off upon puromycin treatment confirms inhibition of translation elongation. **C**. Quantification of mRNA co-localization in untreated versus puromycin-treated cells. Boxplots show the percentage of co-localized transcripts per cell for the indicated mRNA pairs (*FAS1*/*FAS2*, *PFK1*/*PFK2*, *GUS1*/*ARC1*, *GUS1*/*MES1*, *TRP2*/*TRP3*, *GCD11*/*SUI2*, *CPA1*/*CPA2*, *SUI2*/*SUI3*). Each dot represents a single cell; black points indicate medians. Statistical significance compared to untreated controls is shown above each pair. **D.** Representative single z-plane smFISH images showing mRNA co-localization in WT cells following puromycin treatment for the complexes quantified in panel C. mRNAs were visualized in two channels (red and green), nuclei were stained with Hoechst (blue). Scale bar = 2 µm.

**Figure 6.**
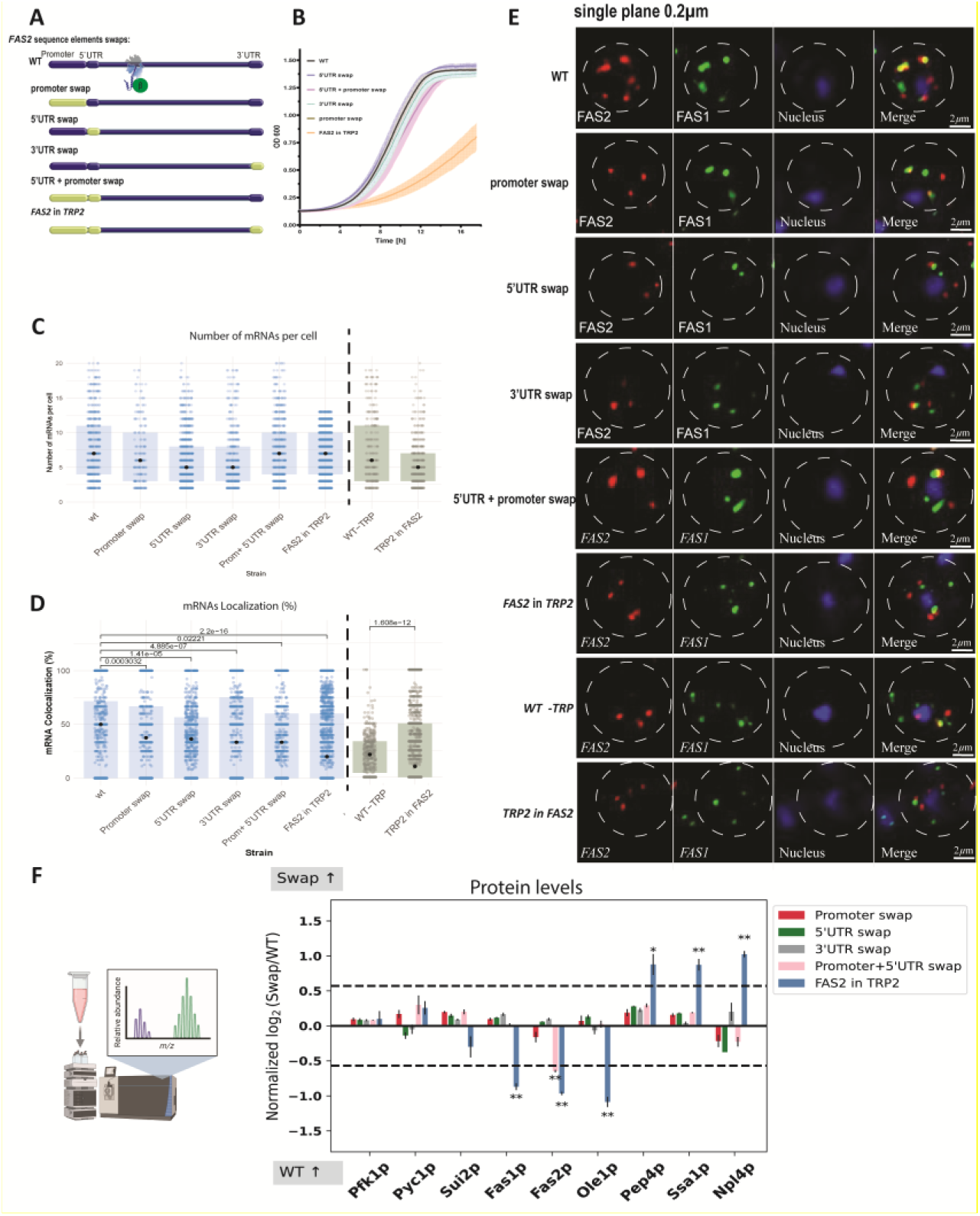
ORF upstream and downstream sequence elements direct the localization of mRNAs encoding for subunits of the FAS complex. **A.** Schematic illustration of the *FAS2* gene, highlighting promoter, 5′UTR, coding region, and 3′UTR elements analyzed. **B.** Quantification of *FAS2* mRNA abundance per cell. Boxplots show the number of mRNA molecules detected by smFISH per cell for the indicated strains. Each dot represents one cell, dot represents the median. **C.** Quantification of mRNA co-localization between FAS1 and FAS2 in the indicated strains (promoter swap, 5′UTR swap, 3′UTR swap, promoter+5′UTR swap, and *FAS2* inserted into the *TRP2* locus). For the reciprocal construct (*TRP2* ORF inserted into the *FAS2* locus), co-localization of *TRP2* and *TRP3* mRNAs was quantified and compared to WT. Boxplots show the percentage of co-localized mRNAs per cell detected by smFISH. Statistical significance was calculated relative to WT; p-values are indicated. Each dot represents one cell (n > 700 cells per strain, ≥ 2 independent experiments). **D.** Proteomic analysis of *FAS2* swap strains. Normalized log₂ protein abundance relative to WT is shown. Bars represent proteins levels of various pathways’ representative, including glycolysis, fatty acid synthesis pathway as well as factors involved in protein quality control. Asterisks indicate statistically significant changes (**p < 0.01). **E.** Representative Z-stacked images of smFISH performed the indicated *FAS2* mutant strains compared to wt. Yeast cells were grown to log phase, formaldehyde-fixed, and hybridized with smFISH probes for co-staining of mRNAs in channel one (in red) and channel two (in green). Nuclei (in blue) were visualized using Hoechst staining (Bar size: 2µm). **F.** yeast growth curve of FAS2 swap strains in fatty acid-deficient medium. OD600 measurements were taken at 10 min intervals over 16 h.

**Figure 7:**
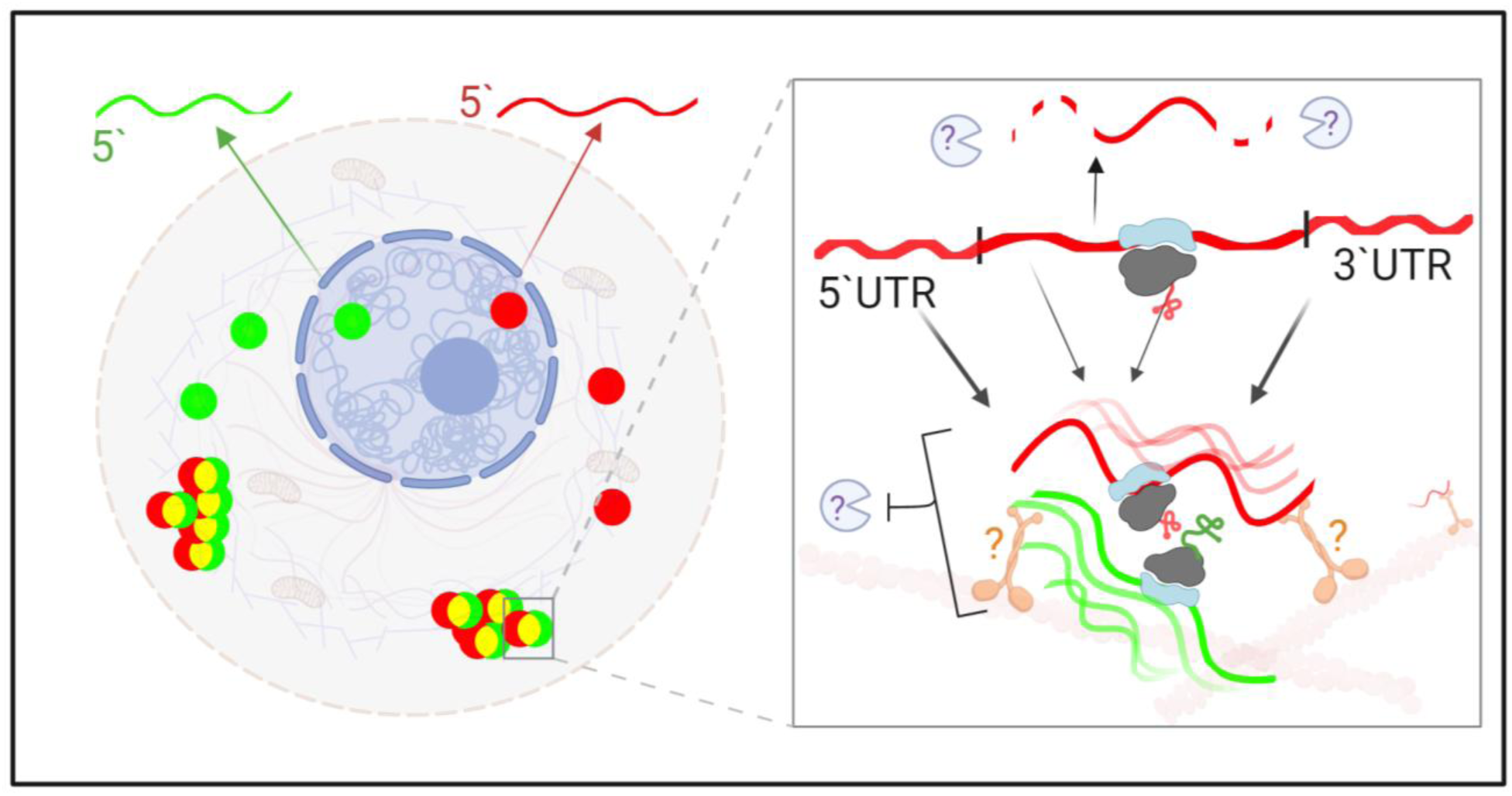
Suggested model of mRNA co-localization mechanisms in Saccharomyces cerevisiae. The left panel illustrates the distribution of mRNAs within the cytoplasm, highlighting the clustering of functionally related mRNAs (green and red) at specific cytoplasmic foci. The right panel provides a detailed view of the structural elements within the mRNAs, including the 5’ and 3’ untranslated regions (UTRs) that contribute to localization. Potential binding factors and molecular interactions are indicated, suggesting their roles in regulating mRNA translation and co-localization, which facilitate efficient protein complex assembly. The question marks indicate areas where further research is needed to clarify the mechanisms involved.

All complexes except CPA displayed significant reduction in mRNA co-localization levels. However, this reduction showed high variability, ranging from ∼25% in *FAS1* and *FAS2* to a stark reduction in *SUI2* and *GCD11* co-localization levels, dropping to a median of 0% (Figure 5C,D). This analysis demonstrated puromycin significantly impacts co-localization, suggesting that nascent chain interactions play a role in mRNA clustering, in a complex-specific manner. However, the results of FAS, PFK, CPA as well as aminoacyl-tRNA synthetase complexes suggest other features are involved in mRNAs clustering.

### 5’ and 3’ UTRs direct co-localization of mRNAs encoding FAS subunits, affecting protein functionality and stability

We investigated whether and which mRNA sequence elements contribute to cytoplasmic foci clustering. Previous studies indicated that mRNA localization often relies on cis-acting elements in their 5’ and/or 3’ UTRs, though coding regions may also play a role (Berkovits & Mayr, 2015; Cassella & Ephrussi, 2022; Chekulaeva, 2024). Additionally, starvation-induced mRNA foci formation is influenced by promoter regions (Chen et al., 2022), suggesting that localization cues may reside in regulatory elements of the genomic locus that are not transcribed. To investigate the impact of these diverse elements on mRNA co-localization in the context of protein complex assembly, we focused on *FAS1* and *FAS2* mRNAs, which encode the hetero-oligomeric FAS complex. We generated yeast strains with the following sequence elements mutated in the *FAS2* genomic locus using CRISPR/Cas9 (Levi & Arava, 2020). as well as SWAp-Tag approaches (Yofe et al., 2016): (1) Promoter swap with *TRP2* corresponding sequences, (2) 5’UTR swap with *TRP2* corresponding sequences (3) Promoter & 5’UTR swap with *TRP2* sequences, (4) *FAS2* ORF swap with *TRP2* ORF, (5) 3’UTR & terminator swap with *CYC1* corresponding sequences (see details in methods).

To assess the impact of *FAS2* mutations on FAS complex formation, we compared growth rates under fatty acid depletion conditions, requiring FAS assembly and function (Figure 5, supplementary Figure 2). Strains lacking essential FAS catalytic domains required fatty acid supplementation for growth, as expected. However, Promoter, 5’UTR, promoter+5’UTR and 3’UTR+terminator swap strains showed growth levels similar to wt (Figure 5B), indicating that these genetic exchanges do not significantly impair FAS complex function, and thus assembly. In contrast, swapping the entire *FAS2* ORF with *TRP2* resulted in an extended lag phase and growth inhibition in fatty acid-deficient media (Figure 5B), suggesting upstream and downstream regulatory elements to the *FAS2* ORF impact FAS assembly and function.

The different cis non-coding element swap strains exhibited *FAS2* transcript levels that were generally comparable to wild type. Strains with the *FAS2* ORF swapped with *TRP2* ORF or those with promoter & 5′UTR swaps had minimal effects on *FAS2* levels, while swaps involving the 3′UTR or 5′UTR resulted in a modest decrease in transcript levels (Figure 5E). We then analyzed the levels of cytosolic co-localization of the *FAS2* and *FAS1* mRNAs (Figure 5D). The distribution pattern of *FAS1* and *FAS2* mRNAs in the wt was ∼50%, while in *FAS2* 5’UTR, 3’UTR, Promoter and Promoter+5’UTR, swap strains showed a modest yet significant reduction compared to wt, to ∼35%. In contrast, swapping the entire *FAS2* ORF resulted in a strong decrease in clustering with *FAS1* to approximately ∼20%. This reduction is even more pronounced, as in this strain *FAS2* mRNA cellular levels remained the same as in wt. Similarly, *TRP2* ORF with *FAS2* resulted in very little impact on *TRP2* mRNA levels yet a significant reduction in co-localization of *TRP2* and *TRP3* levels. Swapping of *FAS2* promoter +5’UTR with the corresponding sequence elements of *NOP1* resulted in comparable results to the swap of these elements with those of *TRP2*: smFISH results showed a slight yet insignificant decrease in co-localization levels and growth assay results showed no significant change compared to wt (supplementary figure 3A, B) This indicates that both upstream and downstream elements in the *FAS2* locus contribute to mRNA co-localization with *FAS1* mRNA, suggesting multiple localization cues contributing in an additive manner. Taken together, these findings suggest that sequence elements in the 5’and 3’UTRs, along with genomic elements further upstream and downstream to the ORF, such as promoter regions, each contribute to *FAS1* and *FAS2* mRNA co-localization. Together, these results suggest a combined role of multiple zip codes with compensatory functions, similar to other mRNA targeting pathways, such as OSKAR and multiple neurites localized transcripts (Gunkel et al., 1998; Mikl et al., 2022). Impairment of these cues compromised FAS complex functionality, highlighting the significance of mRNA localization for complex assembly pathways.

Even subtle changes in translation can have a profound impact, changing the protein’s structure, stability or cellular levels (Komar et al., 2024; Y. Liu et al., 2016). Furthermore, previous analysis by Shiber et al., (2018) demonstrated that deletion of the *FAS1* or *FAS2* genes’ interaction domains (Figure 1C and extended data figure 1 C,D) reduced dramatically co-translational assembly interactions and functionality. Yet, no significant impact on either genès rate of translation was observed by ribosome profiling. Still, it is possible that the 5’ and 3’ cis-elements swaps impact translation efficiency, thus indirectly affecting mRNA localization. Therefore, we tested the impact of swapping of the cis genetic elements on the final product - the protein levels. To capture the impact of the *FAS2* swap strains on protein levels, with high sensitivity and precision, we performed mass-spectrometry based proteomics analysis with isotopic labeling. *FAS2*-swap strains primary amines labelling was performed with light isotope 12-CH2-formaldehyde. Wild type samples were labelled with heavy isotope 13-CD2-formaldehyde and spiked into each swap strain for internal quantification. This was performed in four biological replicates per strain. The results allowed us to test the relative protein abundance changes not only on Fas2 protein levels but also in a global manner. Our results demonstrate there was no significant impact on Fas2 protein levels in these strains: Promoter, 5’UTR or 3’UTR swap strains (Figure 5). In these three strains, where *FAS2* gene cis elements were swapped with *TRP2* gene elements, we did not detect a significant impact on the majority of the proteome. Furthermore, in the strain where both Promoter & 5’UTR of *FAS2* were swapped with those of *TRP2*, we detected only a small, yet reproducible ∼30% decrease in Fas2 protein levels. Altogether, we did not detect a significant impact on the majority of the proteome (figure 5). In the strain where *FAS2* ORF was swapped with *TRP2* ORF, we detected the most significant impact. Fas2 protein levels decreased by 2-fold. Interestingly, Fas1 protein levels also decreased by almost 2-fold. Acc1, which precedes the FAS complex in the fatty acid synthesis pathway, showed slight to no change. Ole1 (OLEic acid requiring stearoyl-CoA 9-desaturase), which functions downstream of the FAS complex (Nguyen et al., 2011) displayed 2-fold reduction. Furthermore, our analysis revealed markedly increased levels of factors targeting mis-assembled FAS complex subunits, suggesting upregulated protein quality control. FAS subunits mis-assembly was shown to trigger targeting to three different cellular pathways: Vacuolar proteolysis by Pep4 (Schüller et al., 1992); Chaperone mediated aggregation by Ssa1, as well as proteasomal degradation, involving the highly conserved Cdc48p-Npl4p-Ufd1p segregate complex (Scazzari et al., 2015). We identified ∼2-fold increase in all these factors: Pep4, Ssa1 and Npl4 (Figure 5).

Taken together, these results suggest that the individual 5’UTR, 3’UTR and promoter gene elements do not significantly affect Fas2 protein levels. This is in accordance with the smFISH results, demonstrating little impact on mRNA levels or co-localization, as well as the growth analysis in condition where FAS activity is essential (fatty acids depletion). Promoter and 5’UTR swap additive impact led to a small yet reproducible decrease in Fas2 protein levels alone. FAS2 ORF swap, combining promoter, 5’UTR and 3’UTR exchange caused the most significant impact on the fatty acid synthesis pathway, leading to reduced protein levels of Fas2, Fas1 and Ole1 of the pathway. Furthermore, increased protein quality control factors targeting misassembled FAS subunits was observed. Thus, the proteome analysis results further support the co-localization analysis, demonstrating the additive impact of cis-acting elements disruption on the downstream protein levels. When mRNA mis-localization occurs, it can impair co-translational interactions, resulting in FAS complex mis assembly, reducing functionality as well as stability. Co-translational interactions can be crucial to achieve correct folding and protect from misfolding, as was recently demonstrated in several biophysical studies. Detailed analysis of isolated ribosome-nascent chain complexes by quantitative NMR as well as Cryo-EM revealed that timely association of co-translational chaperones as well as interactions with the ribosome exit tunnel and surface facilitates sampling of native protein conformations (For example: Galmozzi et al., 2025; Plessa et al., 2021; Roeselová et al., 2024; Streit et al., 2024; Venezian et al., 2024). The results indicate that the spatial distribution of *FAS2* transcripts plays a crucial role in determining the stability of fatty acid synthesis enzymes.

## Discussion

mRNA localization regulation has been implicated in various functions, including embryonic development, proliferation, stress responses, nutrient sensing, and subcellular compartmentalization (Buchan & Parker, 2009; Cohen et al., 2022; Eliyahu et al., 2012; Gadir et al., 2011; Gagnon & Mowry, 2011; Haimovich et al., 2016; Horste et al., 2023; Hoyle & Ashe, 2008; Johnstone & Lasko, 2003; Lehmann’ & Niisslein-Volhard’, 1986; Panasenko et al., 2019; Pizzinga et al., 2019; Balagopal & Parker, 2009; Morales-Polanco et al., 2021). Furthermore, the assembly of several large complexes, composed of dozens of subunits - the nuclear pore complex, RNA polymerase II and III, and the SAGA transcriptional coactivator-was corelated to the spatial organization of mRNAs encoding their subcomplexes. Some transcripts encoding subunits of these assemblies’ exhibit marked cytoplasmic clustering, whereas others do not (Bernardini & Tora, 2024; Hampoelz et al., 2019; Kamenova et al., 2019; Lautier et al., 2021; Mallik et al., 2025; Yayli et al., 2023).

These observations raise several key questions: How abundant is spatial transcripts organization across various complexes? Cellular pathways? How does it impact protein assembly, function, or stability? What features regulate the different patterns of transcripts’ distribution?

The results presented here demonstrate that spatial organization in cytoplasmic foci is an abundant phenomenon among cytosolic complexes in *S. cerevisiae*. This was observed for complexes as small as dimers or trimers (for example the aminoacyl-tRNA synthetases complex) or as large as 12 subunits complex (such as the FAS complex). This pattern was observed across a range of complexes with diverse cellular functions, structural architectures, and regulatory characteristics. It was also found in multiple metabolic pathways, including those involved in fatty acid synthesis, tryptophan biosynthesis, arginine biosynthesis, and the gluconeogenesis-glycolysis regulatory network.

Function-related mRNA co-localization can facilitate early subunit interactions, as soon as interface domains are exposed at the ribosome exit tunnel. In fact, ribosome profiling experiments revealed that partner subunit association indeed occurs right upon the exposure of interaction domains (Shiber et al., 2018; Venezian et al., 2024). Translational pausing was suggested to aid these timely interactions. In *S. cerevisiae*, 78% of transcripts display transient translation slowdown at specific sites, mediated for example by local enrichment in nonoptimal codons (Sharma et al., 2021). Such specific slowdowns are correlated with the co-translational association of the Signal Recognition Particle (SRP), which promotes the docking of ribosome-nascent chain-SRP complexes to the ER membrane translocon (Mason et al., 2000; Pechmann et al., 2014; Voorhees & Hegde, 2015). However, co-translational assembly pathways of heteromeric complexes often do not involve a slowdown in translation at the onset of interactions (Shiber et al., 2018). Our results propose an alternative mechanism in which co-localized translation reduces the need for pausing by increasing the local enrichment of ribosomes synthesizing interaction partners, enhancing association rates. By enabling timely subunit interaction, co-localized mRNA synthesis can protect emerging polypeptide chains from harmful hydrophobic interactions, which can lead to misfolding and aggregation.

Different mRNA features can direct cytoplasmic localization. Zip codes in 5’and 3’UTRs, and in coding regions, were shown to recruit dedicated localization factors as well as molecular motors (Jambhekar & DeRisi, 2007; Kislauskis & Singer, 1992; Martin & Ephrussi, 2009; Mendonsa et al., 2023). Furthermore, stress-induced mRNA distribution in the cytoplasm was shown to depend on promoter regions (Zid & O’Shea, 2014), suggesting transcription factors could additionally function in cytoplasmic targeting. We analyzed the features of *FAS2* mRNA influencing its cellular distribution, finding that swapping its ORF into the *TRP2* locus strongly reduced co-localization with *FAS1* mRNA and impaired growth, suggesting that co-localization is critical for assembling metabolic complexes. Therefore, our results support the hypothesis that mRNA clusters are vital for the assembly of key metabolic complexes. The upstream and downstream untranslated sequences of the ORF are essential for proper cytoplasmic localization, indicating that multiple zip codes may have evolved to ensue, in an additive as well as redundant manner, co-localization with functionally related mRNAs encoding for partner subunits. Further studies are required to elucidate which cellular factors are involved in the cytoplasmic spatial organization, and weather they inhibit translation before co-localization can be achieved. As *FAS2* encoded protein is highly dependent on its partner to achieve correct folding and avoid rapid degradation (Scazzari et al., 2015), co-localized synthesis and assembly may be crucial for the stability of the *FAS2*-encoded protein. Accordingly, disruption of mRNA co-localization levels of transcripts encoding subunits of the fatty acid synthase complex lead to a significant reduction of Fas2 protein levels, concomitantly displaying an elevation in various pathways implicated in the elimination of this subunit. These results strongly indicate that localized synthesis provides early protection from misfolding, in coordination with the translation process. Furthermore, the stability of several enzymes in the fatty acid pathway was impacted, reducing the protein levels of Fas1 and Ole1 for example, while displaying an elevation in distinct pathways involved in the quality control of these proteins, such as Pep4, vacuolar aspartyl protease, similar to lysosomal elimination pathways. These results suggest broader regulatory effects of disruption of mRNAs clustering hubs on complex partners and downstream enzymes in the pathway, which can lead to overwhelming of the cellular quality control pathways, as was observed in aging models (Labbadia & Morimoto, 2015).

Finally, the emerging polypeptide chains may also play a key role in mediating mRNA co-localization in complex specific translation hubs, as evidenced by our finding that puromycin decreases mRNA co-localization levels of several complexes. Puromycin had a variable impact, as was recently demonstrated also by Mallik et al (Mallik et al., 2025). Further corroborating the intricate interplay of multiple targeting cues in spatial organization. Further research will clarify if this is mediated via direct nascent chain interactions, mRNA interactions or whether these interactions require additional factors. For example, a recent study identified a novel mRNA targeting mechanism, involving nascent chain recruitment of molecular motors, as well as mRNA motifs involvement (Cassella & Ephrussi, 2022). We further found widespread mRNA co-localization in key metabolic pathways, including arginine synthesis, glycolysis and fatty acid synthesis. Translational repression of fatty acid synthesis was recently shown to regulate cell survival under glucose starvation in glioblastoma (Levy et al., 2024). Furthermore, metabolic homeostasis regulation by mRNA translation were shown to alleviate effects of excess glucose and fatty acids in tissues such as pancreatic islets and the liver (Schaffer, 2020). Glycolytic enzymes cluster in membrane-less cytoplasmic granules in response to hypoxia (Jin et al. 2017) that rely on the participation of their encoding mRNAs (Fuller et al. 2020). This clustering was suggested to facilitate rapid feedback regulation of the entire pathway translation. Our results suggest that even under physiological conditions, mRNAs clustering occurs, allowing for rapid regulation of fluctuating metabolic needs.

In conclusion, this study indicates that mRNA co-localization is a crucial regulatory mechanism for efficient protein complex assembly, ensuring proper folding and function. This process may significantly impact cellular organization and offer new insights into treating diseases linked to protein misfolding and metabolic dysfunction.

## Acknowledgements

We would like to thank all the Shiber lab members for fruitful discussions, and especially Muhammad Makhzumy for his contribution to co-localization analysis. We thank the Technion’s Lorry I. Lokey Interdisciplinary Center for Life Sciences and Engineering for the core facility. We are grateful for the professional services provided by Nitsan Dahan and Yael Lupu-Haber from the Imaging and BioAnalysis unit of the LS&E Infrastructure Center of the Technion, Israel. This work was funded by the European Research Council Starting Grant 2031817 (A.S.) and the Israeli Science Foundation grants 2106/20 (Q.S.). B.B. acknowledges a research grant of the European Union (ERC - SyG - 101072047 - CoTransComplex). Views and opinions expressed are however those of the authors only and do not necessarily reflect those of the European Union or the European Research Council. Neither the European Union nor the granting authority can be held responsible for them.

## Methods

### Yeast strain constructs

Deletion and various sequence swap strains in the endogenous genomic locus of the *FAS2* gene were generated using homologous recombination, or CRISPR/Cas9 (detailed below) following the method described by Janke (Janke et al., 2004) utilizing SWAT (SWAp-Tag) library strains (Meurer et al., 2018; Weill et al., 2018; Yofe et al., 2016). All experiments were performed in the *S. cerevisiae* BY4741 strain background. *S. cerevisiae.* The primers used for constructing these strains are provided in Supplementary Table 4. The strains used in this study are listed in Supplementary Table 5.

### CRISPR\Cas9

To generate the swapped (mutant) strains used in this study, we used CRISPR/Cas9 genome editing as described by Ofri Levi *et al*. (Levi & Arava, 2020), (Venezian et al., 2024).

### Growth assays

Yeast cultures were incubated in YPD growth medium, or in YPD supplemented with fatty acids (0.03% myristic acid and 1% Tween 40) to test fatty acid auxotrophy levels, as in (Scazzari et al., 2015).

### mRNA smFISH

Yeast cells were grown in SD medium (Low-Fluorescence yeast nitrogen base, Formedium) containing 2% glucose to early log phase growth (to ∼ OD= 0.4); amino acids composition is indicated per experiment. Growth temperature was 30 °C, unless otherwise indicated. The cells were then fixed with 37% formaldehyde for 15 min, centrifuged at 1,200 g for 12 min and re-suspended in 4% paraformaldehyde and 100 mM KPO4 at room temperature for 1 h. After fixation, the cells were sedimented and the pellet was washed then resuspended in 250 µl Zymolyase 20T (2 ml Wash Buffer, 1.8 µl β-mercaptoethanol, 10mM Vanadyl-ribonucleoside complex, 5 mg Zymolyase 20T). Cells were incubated 40 min at 30 °C on shaker, then the cells were sedimented, washed and resuspended with 1ml Triton 100X (0.2% triton in 2% BSA in PBS) and incubated 5 min at room temperature then the cells were washed and incubated overnight in 70% ETOH 1ml at 4 °C. The cells were then washed and incubated in hybridization buffer (4xSSC, 50% formamide, 10% dextrane sulfate, 125 µg/ µl E. coli tRNA, 500 µg/ml salmon sperm DNA, 1x Denhardts solution, 1mM DDT and 0.4 U/µl rRnasin) 1hr at 37 °C. thereafter added smFISH probes for labeling. The sequence-specific fluorescent probes (PixelBiotech GmbH) were generated targeting the Open Reading frame (ORF) of the transcript of interest (see Table 3) unless otherwise indicated. To analyze the impact of these gene modifications on mRNA localization, we used smFISH. Probes targeting the last ∼600 nt of the *FAS2* mRNA coding region were used for most strains, except for the *FAS2* MPTΔ strain (3’UTR probes) and the MPT alone strain (MPT-specific probes). For each transcript, 32 complementary DNA probes were produced (∼17-21 nt in length), each conjugated with 3 fluorophores in tandem (Atto-647 or Atto-565), using a T4 DNA ligase (T4DL) so each mRNA molecule is tagged with 96 fluorophores (Cheng et al., 2017). The nuclear staining was performed using Hoechst (20 ng/µl). The stained cells were resuspended in Mounting medium (55% Glycerol in 1 X SSC and 0.125 M EDTA).

### Confocal Microscopy

High-sensitivity confocal imaging was performed using Spinning Disc Confocal microscope from Nikon with CSU-W1 Confocal Scanner Unit equipped with dual sCMOS cameras from Yokogawa at the LS&E Microscopy Core Facility. Cells were imaged with a magnification 100x Oil Objective and high numerical aperture (NA-1.45), fluorescent filters for DAPI, FITC, Atto-647N and Atto-565.

### mRNA image analysis

Images were first processed using the embedded Nikon NIS-Elements deconvolution software, which employs an embedded software algorithm to mathematically correct image blur caused by the optical system. Each fluorescence channel was analyzed independently by applying a channel-specific intensity threshold, determined by plotting the number of detected spots as a function of the threshold (level of fluorescence when no probes were added as well as deletion strains where fluorescent probes were added where no targets are available) and choosing the value at the curve’s inflection point, to effectively distinguish true mRNA signals from background noise. Spots were defined as three-dimensional connected components within the z-stacks, and only those within a specified minimum and maximum volume were retained to filter out noise. Co-localization was then assessed by calculating the pairwise Euclidean distances between spot centroids in the two channels; spots were considered co-localized if the distance was below a three-voxel threshold, with each spot paired to at most one in the opposite channel to prevent over-counting, using IMARIS software platform for co-localization analysis in three-dimensional fluorescence microscopy datasets.

The quantification is based on the segmentation of the cell (bright field) and nucleus (405 nm) to identify the cell compartment of the mRNA signal, each mRNA foci detection in two channels: detection at 641 nm, detection at 565 nm. A 3D Gaussian mask is used to identify the centroid for each detected dot and the distance between neighboring dots is estimated centroid to centroid. The mRNA quantification per foci is also estimated in this analysis using accumulative intensity Quantification of mRNA count per foci was performed based on: (Zenklusen et al., 2008); comparing dots of various intensity across cells. Validation was performed by comparing dot intensities using FAS1 mRNA probes labelled with either 48 or 96 fluorescent Atto dyes per probe.

### Puromycin treatment

For translation inhibition experiments, BY4741 WT yeasts were grown overnight in YPD medium and subsequently diluted 1:20 into fresh media, followed by incubation for 2.5 h at 30 °C to reach early log phase. The media was then supplemented with SDS (0.003%) and proline (0.1%) as the main nitrogen source, to increase the permeability of the cell wall and membrane to the drug treatment and incubated for 30 min at 30 °C, as described in detail in (C. Liu et al., 2007). Puromycin (1 µg/ml final concentration; Sigma-Aldrich) was added, and cells were further incubated for 60 min at 30 °C before processing for polysome or smFISH analysis as described above.

### polysome profiling

Polysome profiling was performed as described in: (Pospísek & Valásek, 2013).

### Cell growth and protein extraction for Mass-spectrometry (MS)

Four biological replicates of each yeast strain were grown overnight in 5ml YPD at 30°C, 220rpm. In the next day, the yeast culture was diluted to OD600 0.1 in YPD and left to grow until reaching OD600 0.6. Cells were briefly pelleted at 3000g for 5min and resuspended in 5 ml synthetically defined media without tryptophan and left to grow for another 15min before pelleting the cells and washing the pellets three times with DDW. Protein extraction was done by resuspending the pellet with lysis buffer containing 1% SDC, 100 mM HEPES (pH 8) and heating for 10 min at 95 °C while shaking at 900 rpm. Protein concentration was measured using BCA Protein Assay Kit (Thermo Fisher Scientific), 50 µg of protein from each sample were transferred to a clean Eppendorf tube and had the volume completed to 50 µl using lysis buffer. Proteins were then reduced with a final concentration of 5 mM DTT at 65 °C, 600 rpm for 30min and alkylated with 12.5 mM final concentration of CAA for 30 min at room temperature in the dark. Sequencing grade Trypsin (Promega) was added at 1:100 (w/w) and incubated at 37 °C overnight. Digestion termination and SDC depletion were done by adding 10 sample-volumes of water-saturated ethyl acetate, along with 1% of the original sample volume of formic acid. The mixture was vortexed extensively for 1 min then centrifuged at 15000 g for 5 min. The lower aqueous phase containing the peptides was carefully transferred into a clean tube. This process was repeated (without adding formic acid the second time), and after the second centrifugation the upper organic layer was discarded, and the peptide solution was dried to completeness in speed vac (CentriVap Benchtop Vacuum Concentrator - Labconco). Peptides were resuspended in 100mM HEPES pH 7 and the primary amines were labelled using 40mM ^12^CH2-formaldehyde (light) for *Fas2*-swap strains or 13CD2-formaldehyde (heavy) for WT and 20mM sodium cyanoborohydride for 1 hour at 37 °C, 600 rpm. Formaldehyde leftovers were quenched with 100mM final concentration of glycine for another hour at 37 °C. Heavy labelled WT peptides were spiked into each *Fas2*-swap strain before acidifying with 1% formic acid and proceeding to MS analysis.

### Mass-spectrometry analysis

Samples were analyzed with EvoSep One HPLC in line with Orbitrap Exploris 480 Mass spectrometer. Appropriate volumes for 1µg of acidified peptides were loaded onto EvoTips (Evpsep) and were separated over an 88-minute gradient according to the manufacturer standard protocol. MS done in DDA mode, Full MS scans acquired in positive ion mode, scanning from 350 to 1200 m/z, at a 120000 resolution, and normalized AGC of 300%. The top 20 precursors with charge state between 2 to 6 were included for MS2, with an isolation window set to 1.3 m/z, and minimum intensity to 8000. Dynamic exclusion duration of 30s with 10 ppm mass tolerance was enabled. Orbitrap resolution for MS2 scans was set to 15000, normalized AGC of 100%, and HCD fragmentation at 27% NCE.

### MS data analysis

Thermo Raw files were analyzed using MaxQuant Version 2.4.2.0 (Cox & Mann, 2008) and searched against Uniport *Saccharomyces cerevisiae* of July 2025 (including 6730 reviewed protein sequences). MaxQuant searches were carried out using tryptic digestion (Trypsin/P) with 2 missed cleavages allowed. Dimethylation labels were configured for light (C2H4, +28.0313 Da) and (^13^C2D4, +34.0633) at peptide N-termini and lysine residues, multiplicity was set to 2 accordingly. The “Re-quantify” feature was enabled to improve quantification coverage. Carbamidomethylation (+57.021) of cysteine was set as fixed, while oxidation (+15.995) of methionine and acetylation (+42.011) of protein N-terminus were set as variable. Default settings were used for other parameters including minimal peptide length of seven amino acids, 1% false discovery rate were applied at the peptide and protein levels. For statistical analysis, MaxQuant normalized heavy-to-light ratio (H/L) were transformed to logarithmic scale, and replicates of each condition were subject to one-sample t-test against null hypothesis population mean of zero (no change). Significance threshold for relative protein quantification was set to (P-value ≤ 0.05) and a fold-change of at-least 1.5 to either side (Log2(FC) ≥ 0.58). Statistical analyses were performed using Python/SciPy library and data visualization was carried out using matplotlib.

## Data Availability

The mass spectrometry proteomics data have been deposited to the ProteomeXchange Consortium via the PRIDE (Perez-Riverol et al., 2025). Project Name: “Proteome-wide characterization of FAS2 genomic elements swap”. Project accession: PXD068196. (for reviewer access username, Password: eBay4nZJy09O)

## Supplementary information

**Figure 1.**
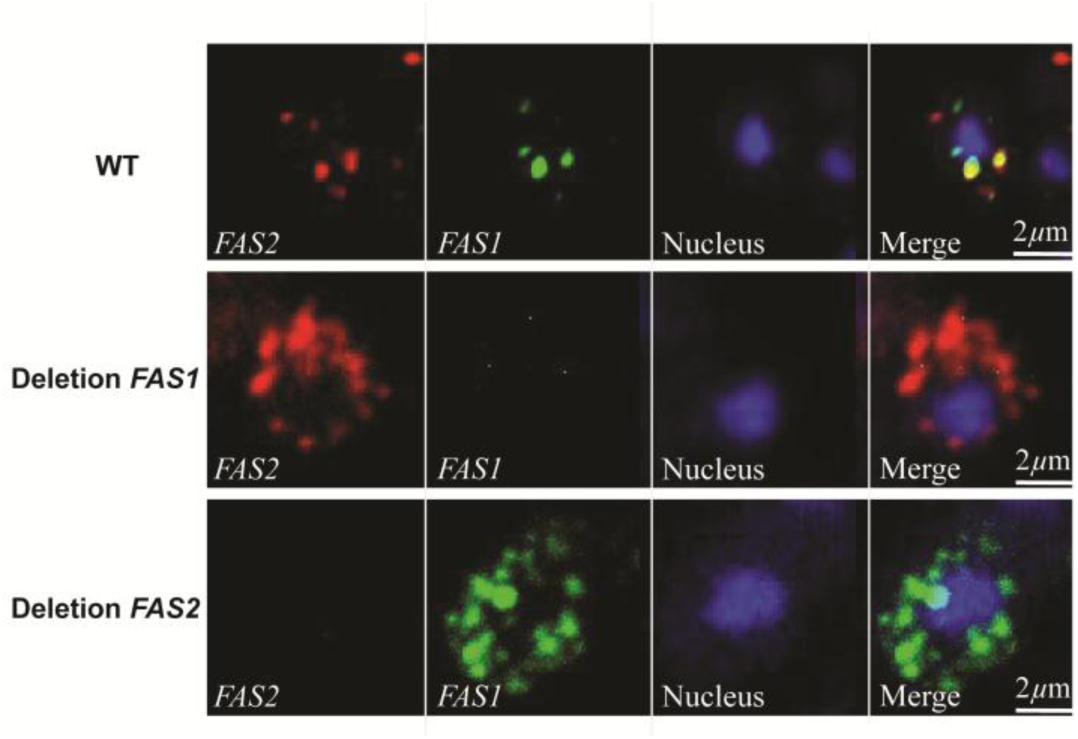
Verification of smFISH probes specificity. Representative Z-stacked images of smFISH performed on wt as well as *FAS1Δ*, and *FAS2Δ* yeast deletion strains for the detection of *FAS1* and *FAS2* mRNAs under physiological conditions. cells were grown to log phase, formaldehyde-fixed, and hybridized with smFISH probes for co-staining of mRNAs in channel one (red) and channel two (green). Nuclei (blue) were visualized using Hoechst staining. Bar size: 5 µm. *FAS2* (red) and *FAS1*(green); mRNAs encoding for subunits of a single protein complex.

**Figure 2.**
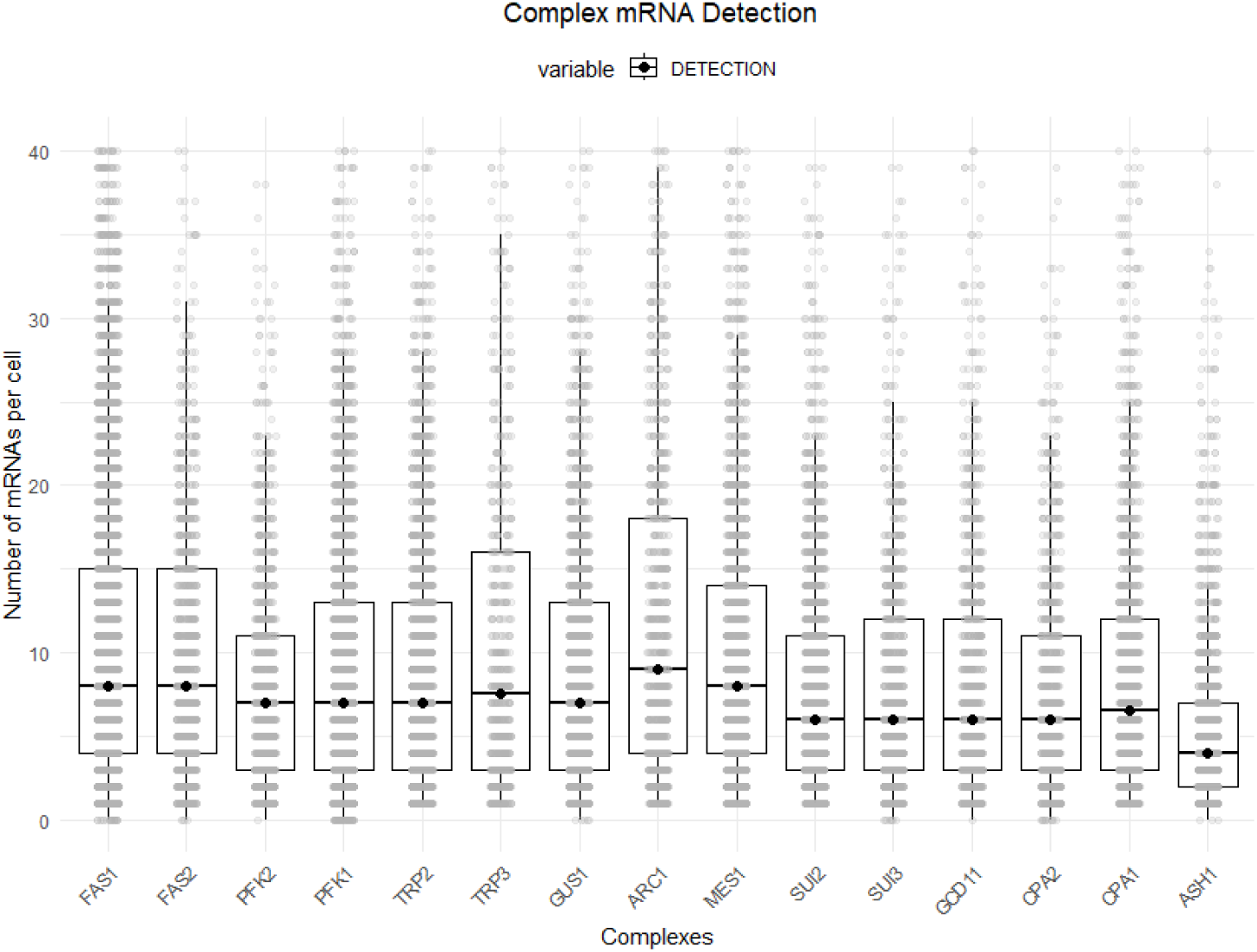
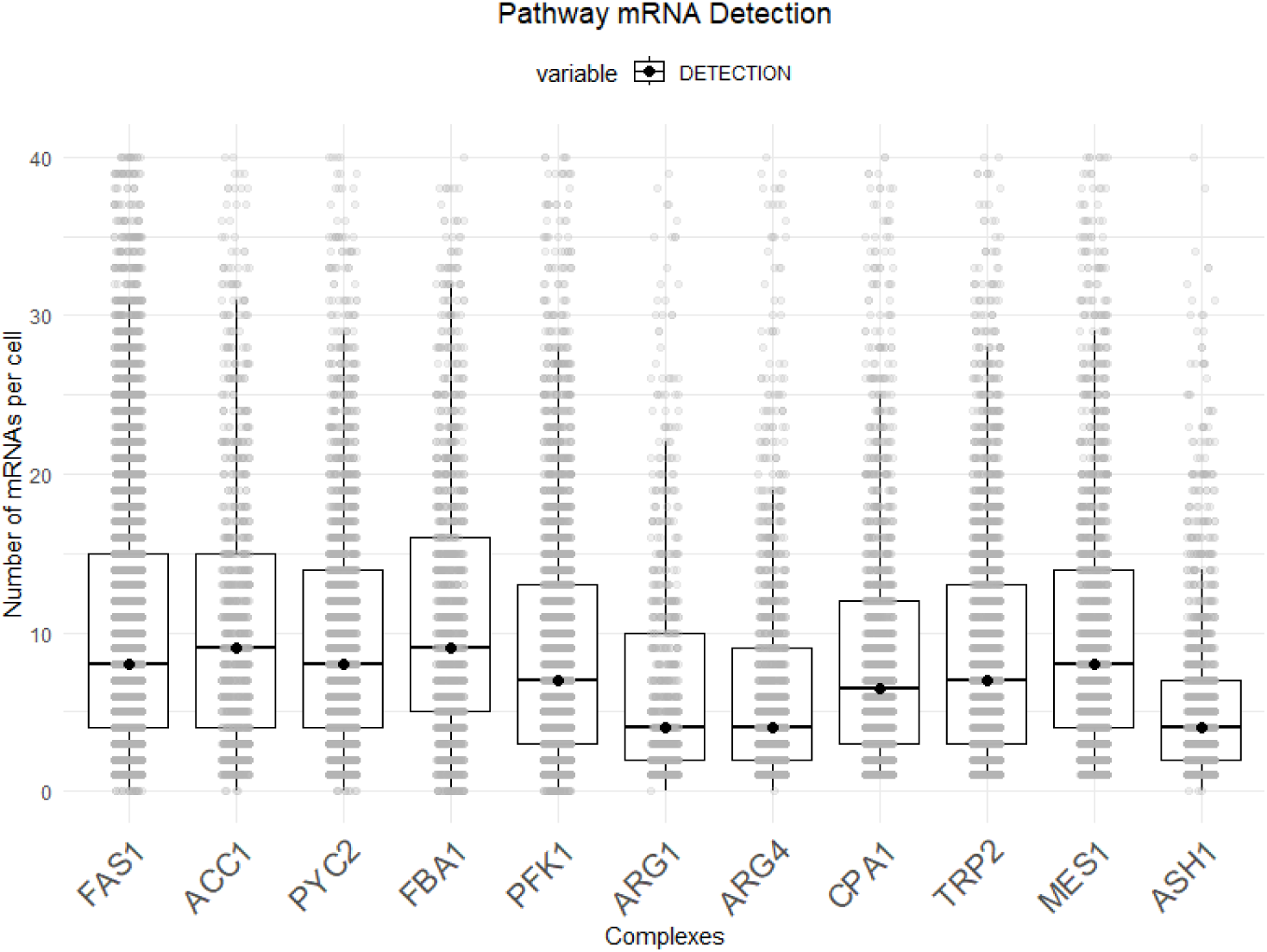
Number of mRNAs per cell detected using smFISH. **A**. Boxplot showing the number of mRNAs per cell for mRNAs encoding for subunits of a single protein complex. **B**. Boxplot showing the number of mRNAs per cell for mRNAs encoding for proteins of a single biosynthetic pathway. Box represents the interquartile range and point represents the median. Each data value represents a single cell, n > 1,319 cells per detection. Two independent experiments.

**Figure 3.**
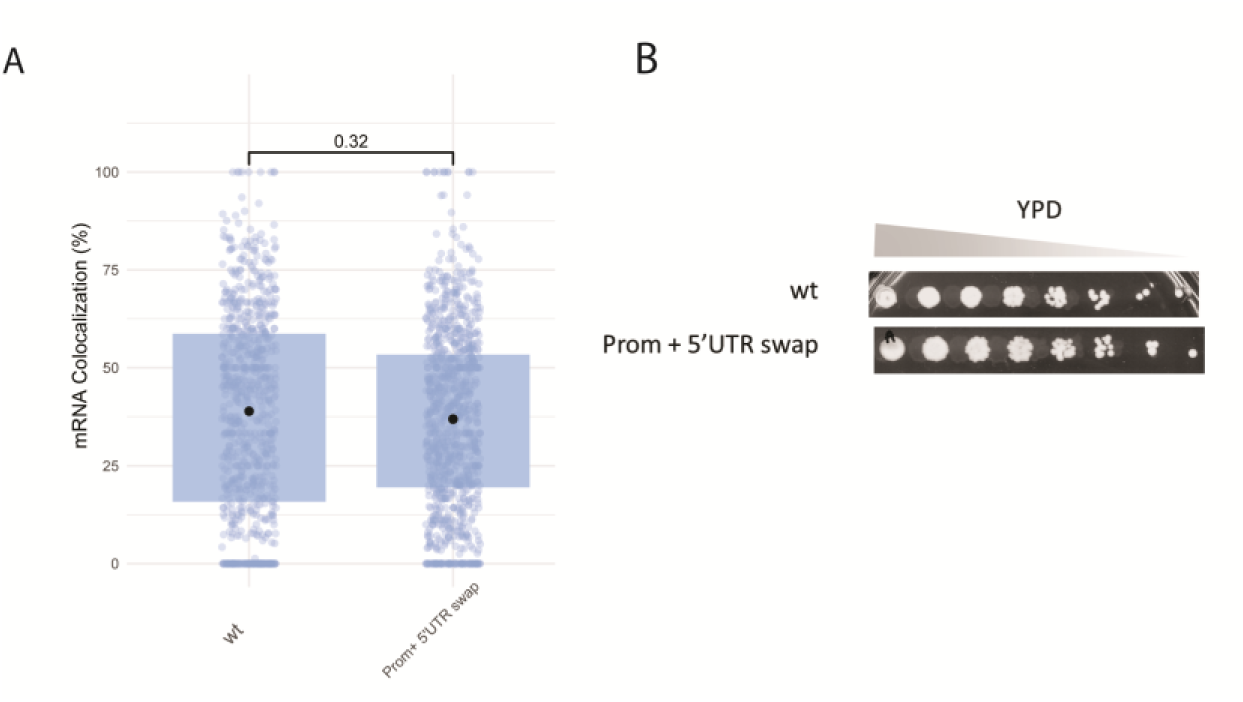
**A**. *FAS2* mRNA co-localization levels following swap with 5’UTR and promoter elements of *NOP1* genomic locus. **B**. Growth assay on YPD plates comparing wild-type (wt) and Prom + 5′UTR *NOP1* swap strains under serial dilution.

**Table 1.**
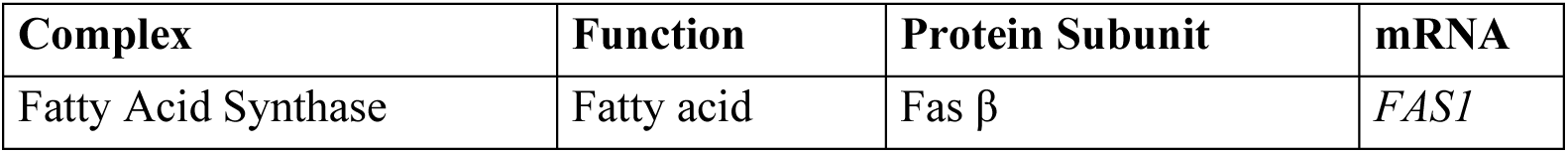
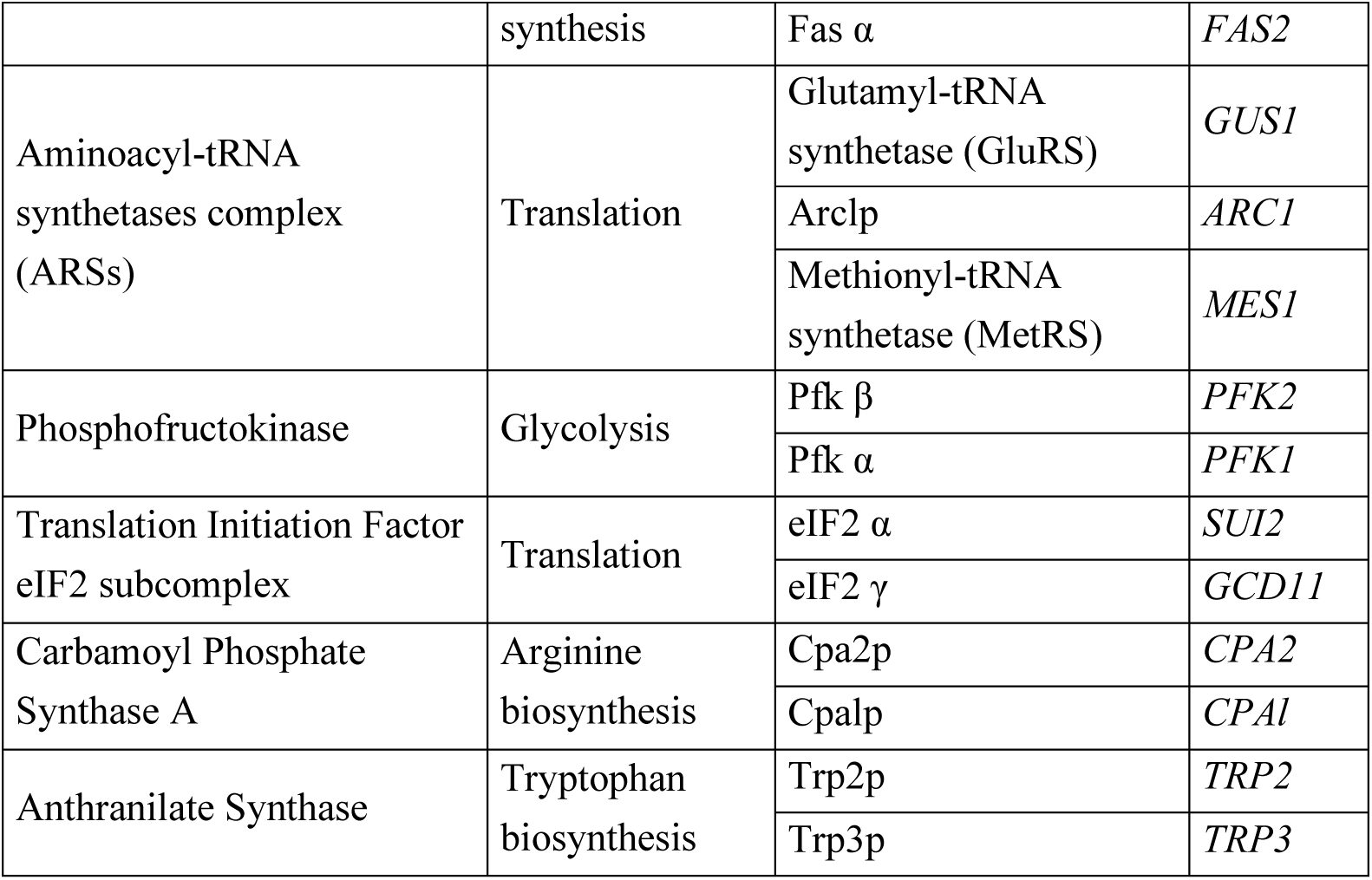
Characteristics of complexes studied, which assemble in a co-translational manner.

**Table 2.** Characteristics of Studied metabolic pathways which transiently interact in a co-translational manner.

| Metabolic Pathway | Protein | mRNA |
| --- | --- | --- |
| Fatty Acid Synthesis | Fas $\beta$ | *FAS 1* |
|  | Acc1 | *ACC1* |
| Arginine biosynthesis | Cpa2p | *CPA2* |
|  | Arg4p | *ARG4* |
|  | Arg1p | *ARG1* |
| Glycolysis & Gluconeogenesis | Pfkβ | *PFK2* |
|  | Pyc2 | *PYC2* |
|  | Fba1 | *FBA1* |

**Table 3.** Chromosomal locations of genes of interest.

| Gene | Chromosome |
| --- | --- |
| *FAS2* | 16 |
| *FAS1* | 11 |
| *PFK2* | 13 |
| *PFK1* | 7 |
| *GUS1* | 7 |
| *ARC1* | 7 |
| *MES1* | 7 |
| *TRP2* | 5 |
| *TRP3* | 11 |
| *SUI2* | 10 |
| *SUI3* | 16 |
| *GCD11* | 5 |
| *CPA1* | 15 |
| *CPA2* | 10 |
| *ACC1* | 14 |
| *PYC2* | 2 |
| *FBA1* | 6 |
| *ARG1* | 15 |
| *ARG4* | 7 |
| *ASH1* | 11 |

**Table 4.**
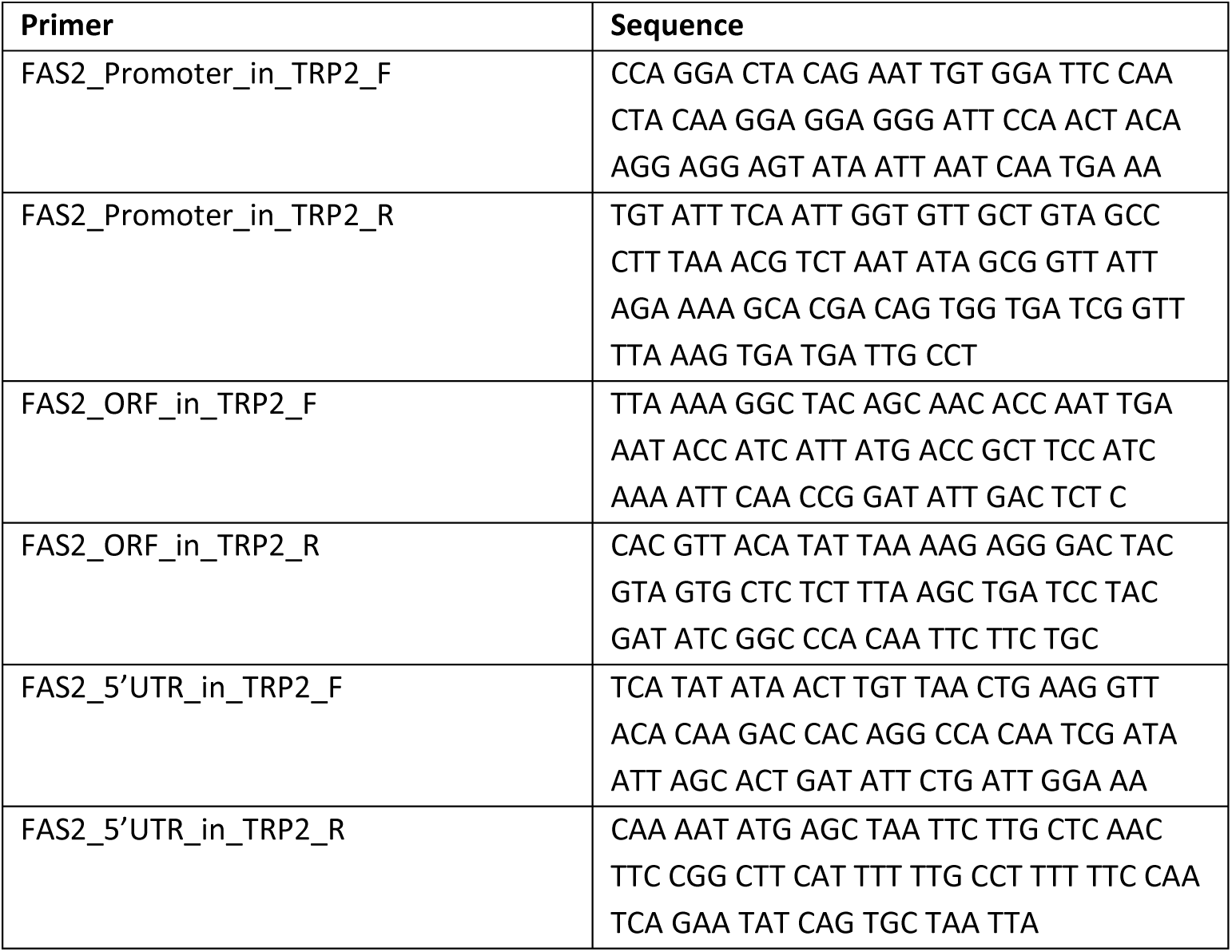
Primers used in this study.

**Table 5.** Yeast strains used in this study.

| Strain | Genotypes | Mating type | Source |
| --- | --- | --- | --- |
| *wt* | his3Δ1, leu2Δ0, met15Δ0, ura3Δ0 | a | EUROSCARF |
| *FAS1Δ* | his3Δ1, leu2Δ0, met15Δ0, ura3Δ0,<br>*FAS1Δ:: His5* | a | This study |
| *FAS2Δ* | his3Δ1, leu2Δ0, met15Δ0, ura3Δ0,<br>*FAS2Δ:: His5* | a | This study |
| SpNOP1pr-sfGFP-FAS2 | his3Δ1 leu2Δ0 met15Δ0 ura3Δ0<br>hphΔn::URA3::SpNOP1pr-sfGFP-FAS2 | a | SWAT LIBRARY |
| FAS2-GFP | his3Δ1 leu2Δ0 met15Δ0 ura3Δ0<br>FAS2-GFP:: hphNT1 | a | Shiber et al.,<br>2018 |
| FAS2 in TRP2 | his3Δ1, leu2Δ0, met15Δ0, ura3Δ0,<br>*FAS2Δ:: His5, FAS2:: TRP2* | a | This study |
| FAS2 MPT domain | his3Δ1 leu2Δ0 met15Δ0 ura3Δ0<br>FAS2 MPT domain (1-282nt)-GFP::<br>hphNT1 | a | This study |
| FAS2 MPT domain in TRP2 | his3Δ1, leu2Δ0, met15Δ0, ura3Δ0,<br>*FAS2Δ:: His5, FAS2 MPT domain (1-282nt)-GFP:: TRP2* | a | This study |

